# Effect of the macroalgae *Asparagopsis taxiformis* on methane production and the rumen microbiome assemblage

**DOI:** 10.1101/436568

**Authors:** Breanna Michelle Roque, Charles Garrett Brooke, Joshua Ladau, Tamsen Polley, Lyndsey Marsh, Negeen Najafi, Pramod Pandey, Latika Singh, Joan King Salwen, Emiley Eloe-Fadrosh, Ermias Kebreab, Matthias Hess

## Abstract

**Background:** Recent studies using batch-fermentation suggest that the red macroalgae *Asparagopsis taxiformis* might reduce methane (CH_4_) emission from beef cattle by up to ~99% when added to rhodes grass hay, a common feed in the Australian beef industry. These experiments have shown significant reductions in methane without compromising other fermentation parameters (i.e. volatile fatty acid production) with *A. taxiformis* organic matter (OM) inclusion rates of up to 5%. In the study presented here, *A. taxiformis* was evaluated for its ability to reduce methane production from dairy cattle fed a mixed ration widely utilized in California; the largest milk producer in the US.

**Results:** Fermentation in a semi-continuous *in-vitro* rumen system suggests that *A. taxiformis* can reduce methane production from enteric fermentation in dairy cattle by 95% when added at a 5% OM inclusion rate without any obvious negative impacts on volatile fatty acid production. High-throughput 16S ribosomal RNA (rRNA) gene amplicon sequencing showed that seaweed amendment effects rumen microbiome communities consistent with the Anna Karenina hypothesis, with increased beta-diversity, over time scales of approximately three days. The relative abundance of methanogens in the fermentation vessels amended with *A. taxiformis* decreased significantly compared to control vessels, but this reduction in methanogen abundance was only significant when averaged over the course of the experiment. Alternatively, significant reductions of methane in the *A. taxiformis* amended vessels was measured in the early stages of the experiment. This suggests that *A. taxiformis* has an immediate effect on the metabolic functionality of rumen methanogens whereas its impact on microbiome assemblage, specifically methanogen abundance, is delayed.

**Conclusions:** The methane reducing effect of *A. taxiformis* during rumen fermentation makes this macroalgae a promising candidate as a biotic methane mitigation strategy in the largest milk producing state in the US. But its effect *in-vivo* (i.e. in dairy cattle) remains to be investigated in animal trials. Furthermore, to obtain a holistic understanding of the biochemistry responsible for the significant reduction of methane, gene expression profiles of the rumen microbiome and the host animal are warranted.

## BACKGROUND

Methane (CH_4_) is a major greenhouse gas with a global warming potential 28-fold greater than that of carbon dioxide (CO_2_) on a 100-year scale [1] and it accounts for approximately 11% of the greenhouse gas (GHG) emissions in the US [2]. Enteric fermentation from ruminant animals alone accounts for approximately 25% of the total CH_4_ emissions in the US, representing the largest anthropogenic source of CH_4_ [3]. Increasing emphasis on reducing GHG emissions from the livestock industry requires advanced methods for reducing and controlling CH_4_ production. Identifying efficient strategies to lower enteric CH_4_ production could result in a significantly reduced carbon footprint from animal production and provide the cattle industry with a way to meet legislative requirements, requiring a reduction of CH_4_ emission of ~40% by 2030.

The biological production of CH_4_ in the rumen is the product of symbiotic relationships between fiber degrading bacteria, hydrogen (H_2_) producing protozoa and methanogenic archaea [4, 5]. Besides being converted into CH_4_, metabolic H_2_ may also be incorporated into volatile fatty acids (VFA), such as acetate, propionate, and butyrate which are then used as energy by the ruminant animal. Theoretically, inhibiting methanogenesis could free molecular hydrogen for use in pathways that produce metabolites (i.e. VFAs) that are more favorable to the host animal, thus creating potential for increased feed efficiency. Since production of enteric CH_4_ can account for up to 12% of the total energy consumed by the animal [6, 7] even a small reduction of CH_4_ production and redirection of carbon molecules into more favorable compounds has the potential to result in significantly more economically and ecologically sustainable production practices in the ruminant industry.

Extensive research has been performed on the effectiveness of feed supplements to reduce enteric CH_4_ emissions through inhibition of microbial methanogenesis within the rumen system [8]. Results have been reported for a number of feed supplements including inhibitors, ionophores, electron receptors, plant bioactive compounds, dietary lipids, exogenous enzymes, and direct-fed microbials indicating reductions on CH_4_ production [9]. While several of these compounds have been shown to inhibit ruminal methanogenesis, some have been shown to decrease VFA production [10] which decreases overall nutrient availability to the animal and is therefore a non-desirable side effect.

Algae are a stable component of the human diet in some cultures [11] and have also been used as feed for agricultural products such as abalone [12] and shrimp [13] The ability of algae to promote well-being and health is mediated to a great extent by highly bioactive secondary metabolites [14, 15, 16] that are synthesized by some of the algal species [17]. In addition to the health promoting properties of macroalgae, some of the brown and red macroalgae have shown to inhibit microbial methanogenesis when tested *in-vitro* [18] and a similar response of the animal microbiome has been proposed. These findings suggest that macroalgae could promote higher growth rates and feed conversion efficiencies in ruminants [19, 20]. Previous findings also suggest macroalgal feed supplements work as highly potent mitigation strategy to reduce CH_4_ production during enteric fermentation [10, 18, 21, 22). Macroalgae feed supplementation therefore may be an effective strategy to simultaneously improve profitability and sustainability of dairy operations.

Various types of algae have antibacterial, antiviral, antioxidant, anti-inflammatory, and anti-carcinogenic properties [23, 24, 25, 26]. Most recently, macroalgae has been tested *in-vitro* and *in-vivo* to determine if there are anti-methanogenic properties within selected types of macroalgae. *Asparagopsis taxiformis*, a red macroalgae, stands out as the most effective species of macroalgae to reduce methane production.

In a recent study [18] *Asparagopsis taxiformis*, a red macroalgae, stood out as the most effective species of macroalgae to reduce methane production. In this work, the effect of a large variety of macroalgal species including freshwater, green, red, and brown algae on CH_4_ production during *in-vitro* incubation was compared. Results showed that *A. taxiformis* amendment yielded the most significant reduction (~98.9%) of CH_4_ production. Moreover, Additional *in-vitro* test with *A. taxiformis* supplementation at inclusion rates up to 5% organic matter (OM) revealed methane reduction by 99%, without significant negative impact on VFA profiles and OM digestibility [10]. Furthermore, this group sought out to identify the anti-methanogenic properties of *A. taxiformis* and found that this particular strain of macroalgae contains an abundance of anti-methanogenic compounds including: bromoform, dibromocholoromethane, bromochloroacetic acid, dibromoacetic acid, and dichloromethane [27]. Bromoform, the most abundant compound found in *A. taxiformis*, was previously identified as a halomethane and has been shown to inhibit enzymatic activities by binding to vitamin B_12_ [28], which chemically resembles coenzyme F430 a cofactor needed for methanogenesis [29]. While it is clear that *A. taxiformis* contains antimethanogenic compounds, actual concentrations of these compounds seem to vary and what causes these variations remain unclear.

In the work presented here, we studied the effect of *A. taxiformis* (5% OM inclusion rate) on the rumen microbiome assemblage and function during *in-vitro* fermentation over the duration of four days. To obtain a better understanding of how this macroalgae would affect CH_4_ emission, specifically from dairy cows fed a diet commonly used California, and therefore providing insight into the value of an *A. taxiformis*-based CH_4_ mitigation strategy for the dairy industry in California. To the end of obtaining new insights into the effect of *A. taxiformis* supplementation on rumen microbiome assemblage, we employed high-throughput 16S rRNA amplicon sequencing. To our knowledge this is the first time that this highly efficient procedure was employed to dissect the changes in the rumen microbiome of dairy cattle in response to *A. taxiformis* as feed supplement and CH_4_ mitigator.

## RESULTS

### *In-vitro* standard measurements remained stable throughout the experiment

Temperature, pH, and mV remained relatively constant (37°C ±2, 6.8 pH ±0.03, 21 mV ±3) throughout the entire experiment and between individual vessels.

### *A. taxiformis* contains an elevated mineral profile but less organic matter compared to SBR

A higher OM content for SBR was found (92.8% DM) when compared to *A. taxiformis* (53% DM). Crude protein amounts were relatively similar for SBR (20% DM) and *A. taxiformis* (17.8% DM). Neutral detergent fiber composition of SBR and *A. taxiformis* were also similar with 38.1 and 36.9% DM, respectively. Differences in starch content between SBR and *A. taxiformis* were prominent with 12.6 and 0.7% DM, respectively. Lignin content for SBR was determined with 6% DM and 4.4% DM for *A. taxiformis*. Total digestible nutrient content (TDN) for *A. taxiformis* was approximately half (33.8% DM) of the TDN determined for SBR (66.2% DM). *Asparagopsis taxiformis* contained elevated mineral profiles compared to SBR. More specifically, *A. taxiformis* exhibited higher calcium, sodium, magnesium, iron, and manganese concentrations. Zinc was present at 23.7 ppm in both SBR and *A. taxiformis*. The detailed composition of SBR and *A. taxiformis* is shown in Table 1.

**Table 1.**
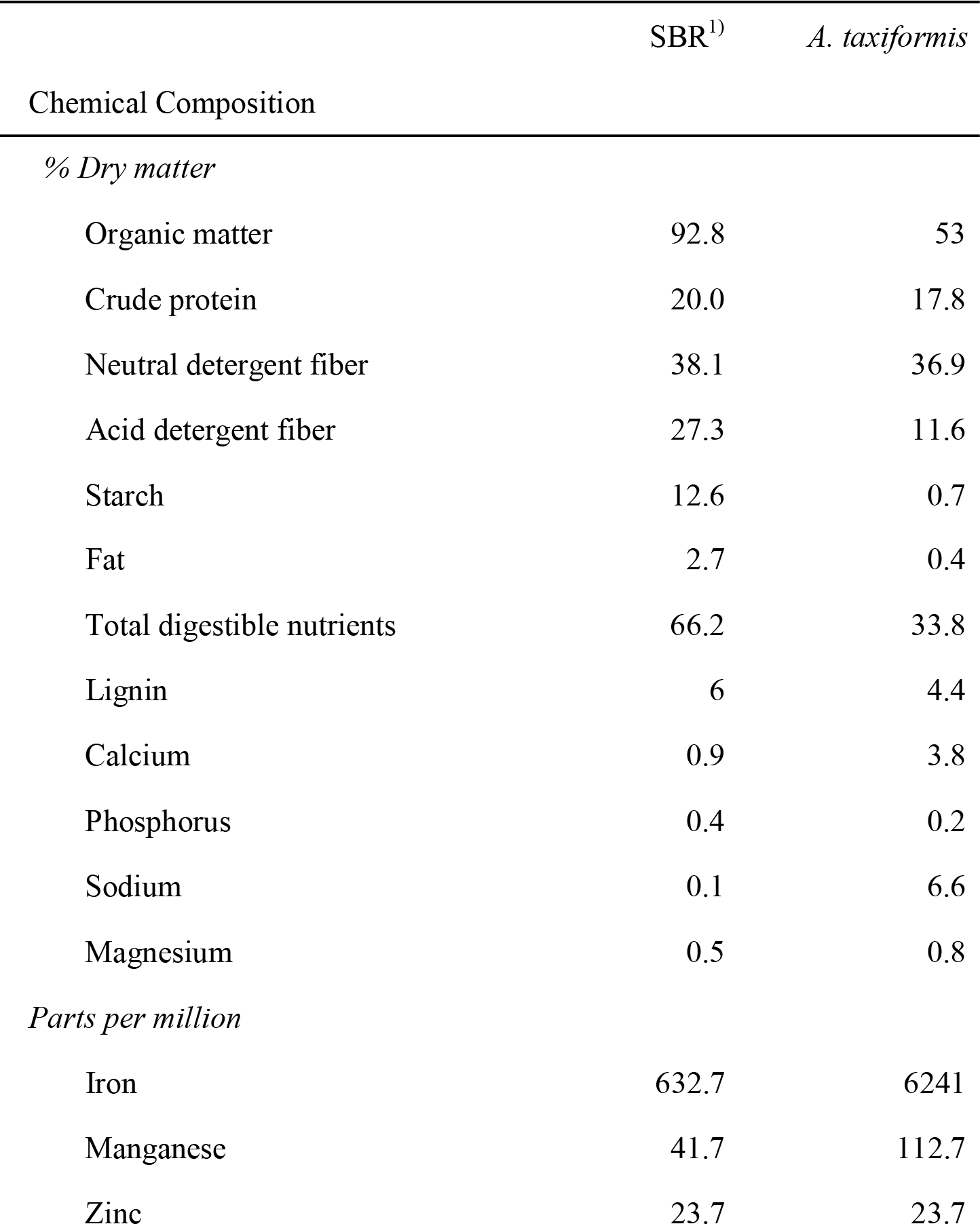
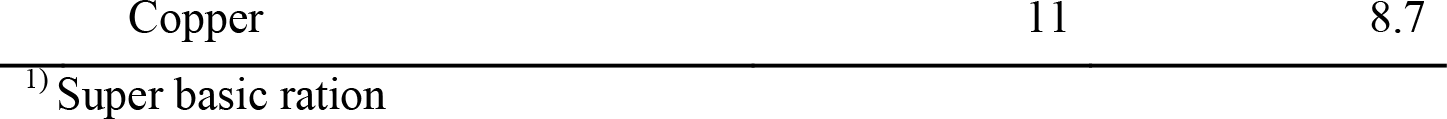
Composition of SBR and *Asparagopsis taxiformis*

|  | SBR<sup>1)</sup> | *A. taxiformis* |
| --- | --- | --- |
| Chemical Composition |  |  |
| % Dry matter |  |  |
| Organic matter | 92.8 | 53 |
| Crude protein | 20.0 | 17.8 |
| Neutral detergent fiber | 38.1 | 36.9 |
| Acid detergent fiber | 27.3 | 11.6 |
| Starch | 12.6 | 0.7 |
| Fat | 2.7 | 0.4 |
| Total digestible nutrients | 66.2 | 33.8 |
| Lignin | 6 | 4.4 |
| Calcium | 0.9 | 3.8 |
| Phosphorus | 0.4 | 0.2 |
| Sodium | 0.1 | 6.6 |
| Magnesium | 0.5 | 0.8 |
| Parts per million |  |  |
| Iron | 632.7 | 6241 |
| Manganese | 41.7 | 112.7 |
| Zinc | 23.7 | 23.7 |

### *A. taxiformis* decreases methane production and increases propionate:acetate ratio

Total gas production (TGP) and CH_4_ production were significantly affected by the inclusion of *A. taxiformis* (*p* < 0.05, Table 2). Average total gas production for the *A. taxiformis* treatment group was 14.81 mL/(g OM) whereas the control group was 28.54 mL/(g OM), representing a 51.8% reduction in TGP with *A. taxiformis*. Average CH_4_ production for the *A. taxiformis* treatment group was 0.59 mg/(g OM), whereas the control group produced 12.08 mg/(g OM), representing a 95% reduction of CH_4_ being synthesized. No significant difference was found in CO_2_ production between the *A. taxiformis* treatment and the control groups. Figure 1 illustrates how total gas (i.e. CH_4_ and CO_2_) was affected over the duration of the experiment. It appears that *A. taxiformis* is effective at reducing TGP and CH_4_ almost immediately, beginning at the 12 hour time point, and continues to inhibit CH_4_ production over 24 hrs just prior to when new bioactive is provided during the feeding process (at 24 hr, 48 hr, and 72 hr). Inhibition of methanogenesis was also measured just prior to the termination of the experiment (96 hr).

**Figure 1.**
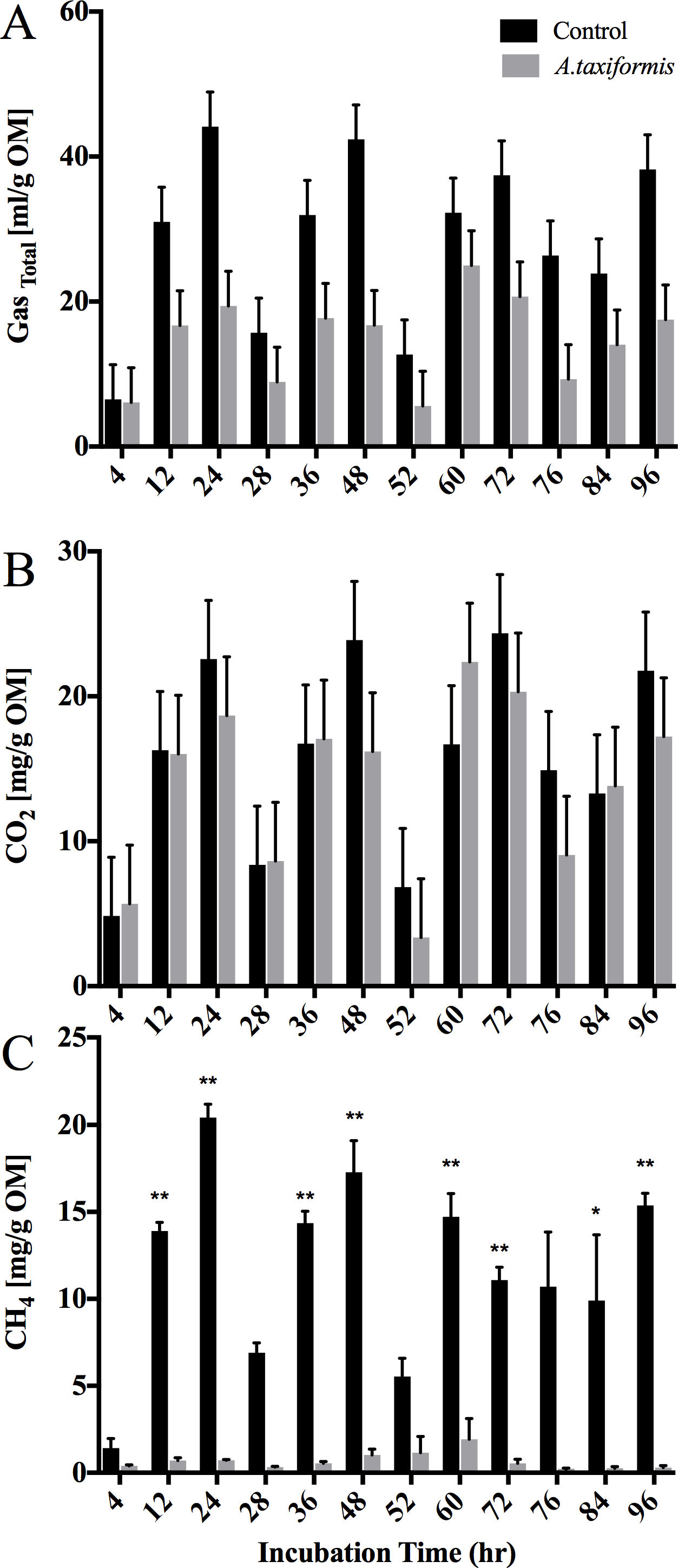
Total gas, CH_4_, and CO_2_ production during *in-vitro* fermentation. Production of total gas [ml/(g OM)], CH_4_ [mg/(g OM)] and CO_2_ [mg/(g OM)] from vessels without (n=3) and with (n=3) *A. taxiformis* as additive at 4, 12, and 24 h over the course of the experiment. **(A)** Total gas production; **(B)** CH_4_ production; **(C)** CO_2_ production. Measurement were performed in triplicates. “**” indicates significant difference (p value ≤ 0.05), “*“ indicates trend toward significance (0.05 > p value ≤ 0.1).

**Table 2.**
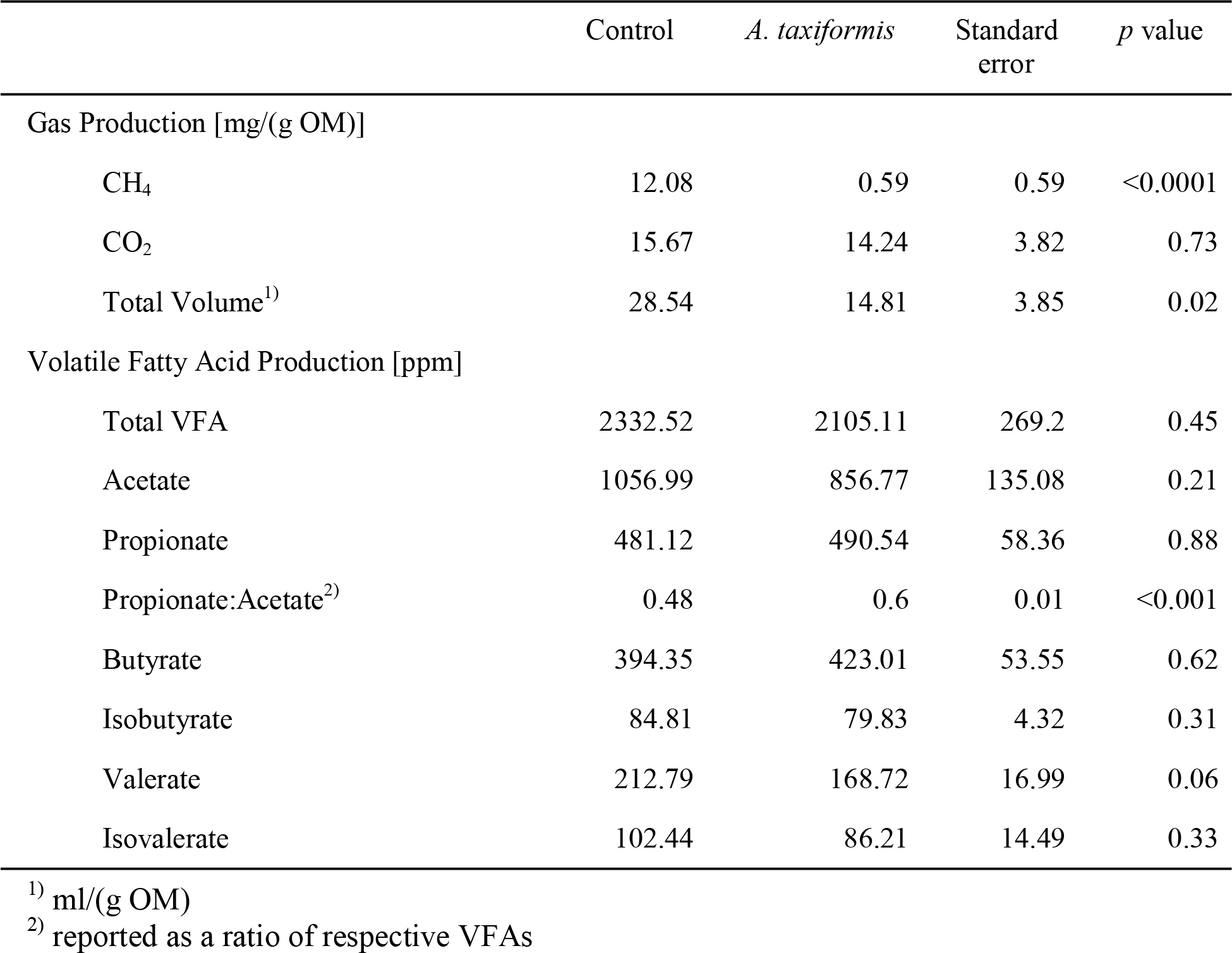
Effects of *A. taxiformis* on total gas production and total volatile fatty acid production.

|  | Control | *A. taxiformis* | Standard error | *p* value |
| --- | --- | --- | --- | --- |
| Gas Production [mg/(g OM)] |  |  |  |  |
| CH<sub>4</sub> | 12.08 | 0.59 | 0.59 | <0.0001 |
| CO<sub>2</sub> | 15.67 | 14.24 | 3.82 | 0.73 |
| Total Volume<sup>1)</sup> | 28.54 | 14.81 | 3.85 | 0.02 |
| Volatile Fatty Acid Production [ppm] |  |  |  |  |
| Total VFA | 2332.52 | 2105.11 | 269.2 | 0.45 |
| Acetate | 1056.99 | 856.77 | 135.08 | 0.21 |
| Propionate | 481.12 | 490.54 | 58.36 | 0.88 |
| Propionate:Acetate<sup>2)</sup> | 0.48 | 0.6 | 0.01 | <0.001 |
| Butyrate | 394.35 | 423.01 | 53.55 | 0.62 |
| Isobutyrate | 84.81 | 79.83 | 4.32 | 0.31 |
| Valerate | 212.79 | 168.72 | 16.99 | 0.06 |
| Isovalerate | 102.44 | 86.21 | 14.49 | 0.33 |

<sup>1)</sup> ml/(g OM)
<sup>2)</sup> reported as a ratio of respective VFAs

Slightly higher total VFA concentrations were recorded for the control group when compared to the *A. taxiformis* treatment group [2332.52 ppm vs. 2105.11 ppm ± 269.20 ppm respectively (means ± SE),] however this difference was not statistically significant (*p* = 0.45, Table 2). Additionally, no significant differences were found when comparing concentrations of acetate, propionate, butyrate, isobutyrate, valerate, and isovalerate (Table 2) between control and *A. taxiformis* treatment group. Although, valerate was not found to be statistically different between groups (*p* < 0.05), it was observed that the *A. taxiformis* treatment group tended to have lowered concentrations of valerate when compared to the control group (*p* = 0.06). Statistical differences were found between groups when comparing the propionate:acetate ratio, with a higher proportion of propionate to acetate within the *A. taxiformis* treatment groups (*p* = 0.001). Differences observed at each timepoint between control and *A. taxiformis* treatment groups were determined to be not significant (Figure 2).

**Figure 2.**
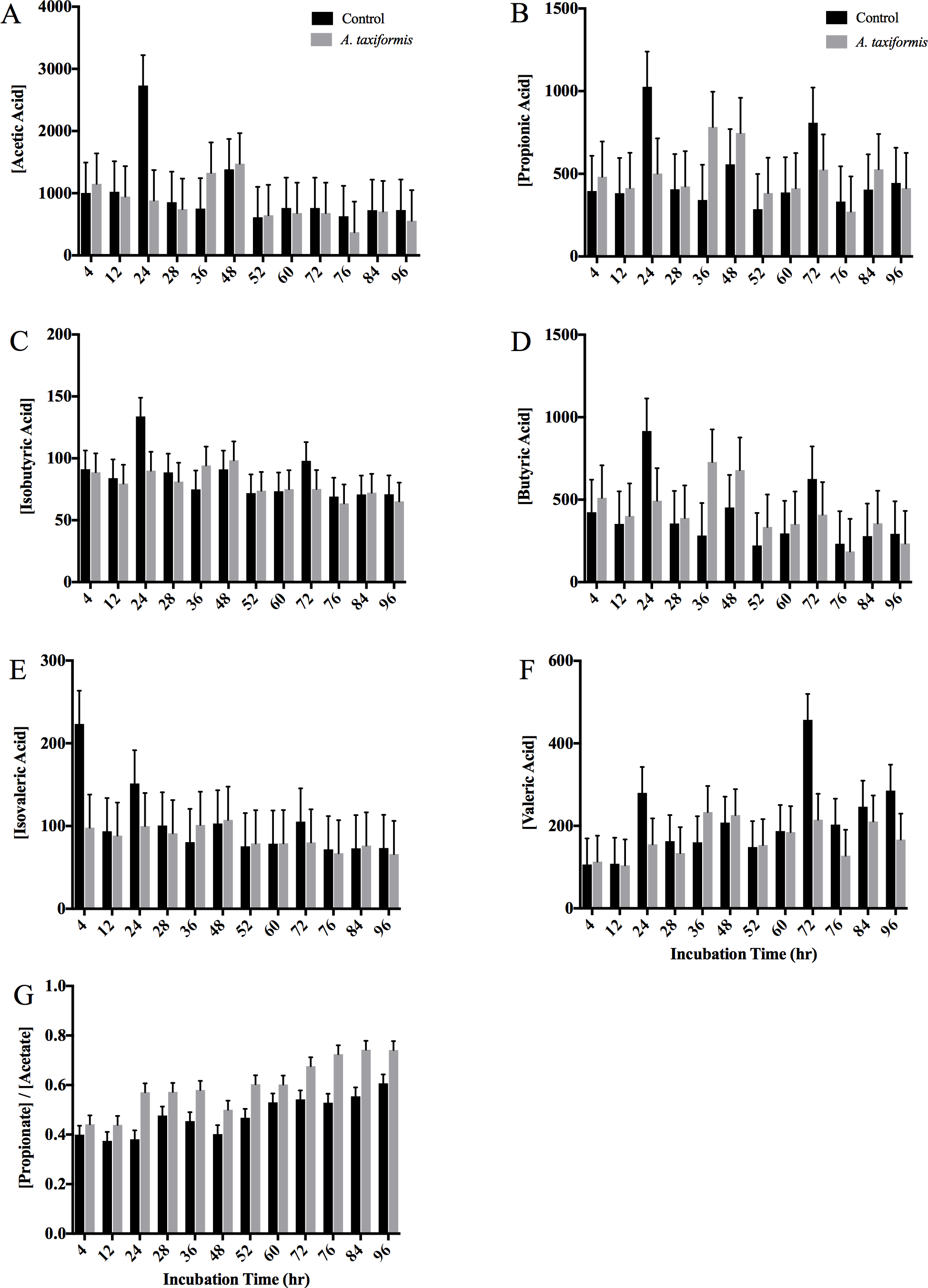
Volatile Fatty acid production during *in-vitro* fermentation. Volatile fatty acid concentrations [ppm] of fermentation fluid of vessels without (n=3) and with (n=3) *A. taxiformis* as additive, determined 4, 12, and 24 h after feeding over 4 days. **(A)** Acetic acid; **(B)** Propionic acid; **(C)** Isobutyric acid; **(D)** Butyric acid; **(E)** Isovaleric acid **(F)** Valeric acid; **(G)** Propionate/Acetate Ratio. Measurement were performed in triplicates.

### Sequencing and quality filtering

A total of 1,251,439 reads were generated from a total of 77 samples, with a mean (± SD) of 16,275 (±1,879) reads per sample. After quality filtering, 757,325 (60.5%) high quality sequences remained. Operational taxonomic units (OTU) based analysis (at 97% sequence identity) revealed 32,225 unique OTUs across all samples. Singletons contributed 23,043 (3%) unique reads to the total filtered read count, and were removed prior further analysis. The mean Goods’ coverage for all samples was 88 ±3%, suggesting that the sequencing effort recovered a large proportion of the microbial diversity in each of the samples under investigation. Distribution of the number of OTUs among each condition and time point during the experiment can be found in Supplementary Table S1.

### α-diversity measurements show microbial communities diverged slightly over the course of the experiment

The microbial communities of the control and *A. taxiformis* amended vessels were compared at each incubation time. Significant differences in the microbial community between the two conditions appeared transiently at only two time points, the 12 hour time point on the first day of the experiment and again at the 24 hour time point on the fourth day (96 hrs after the start of the experiment, AMOVA, *p* ≤ 0.02, and *p* ≤ 0.04 respectively). Comparison of the microbial communities from the start and end of the experiment within each group suggested that the microbial communities changed over the course of the experiment (AMOVA, *p* ≤ 0.06 and *p* ≤ 0.05, treatment and control respectively). The divergence of the microbial communities throughout the experiment was visualized by Principal Coordinate Analysis (PCoA) and is illustrated in Supplemental Figure S2. The first two axes of the PCoA plot account for a low amount of the total variation among the samples (13.5%), which coincides with finding that the two vessel groups were largely similar.

### Microbial communities respond to *A. taxiformis* as a stressor, but recover quickly

Although the effects of seaweed amendments on methane production were immediate (≤ 12 hr), amendments may also affect microbial populations on a longer time scale. Over the duration of the experiment, the β-diversity (turnover) between pairs of control vessels remained constant, but the β-diversity between pairs of treatment vessels and between treatment and control vessels was variable: β-diversity amongst treatment vessels increased and then decreased, peaking at near 72 hours, while β-diversity between treatment and control vessels increased essentially monotonically until the end of the experiment (Figure 3A). These slow shifts in community composition were evident regardless of the taxonomic level at which beta-diversity was considered, including at coarse taxonomic resolutions (Figure 3B). Examination of the genus-level β-diversity within vessels across different time lags also indicated that the microbial communities continued to shift throughout the duration of the experiment (Figure 3C).

**Figure 3.**
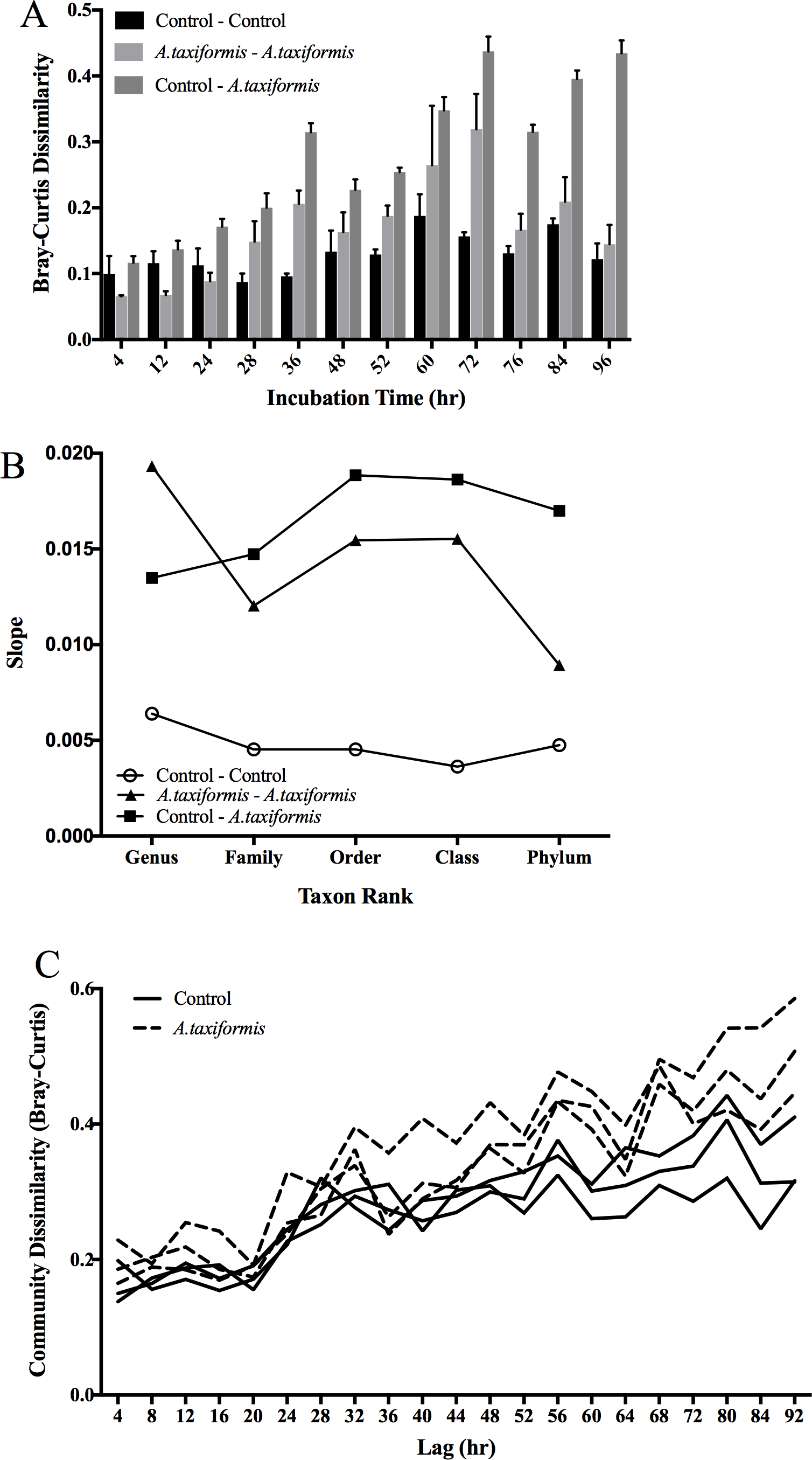
Long-term effects of seaweed amendments on *in-vitro* rumen microbial community. (**A**) Genus-level β-diversity between pairs of vessels at parallel incubation times. (**B**) β-diversity across multiple taxon ranks measured by the slope of the regression of beta-diversity on time for each of the 6 vessels. (**C**) Genus-level β-diversity within vessels at pairs of sampling times.

### Average methanogen abundance decreased, but not in concert with methane reduction

Across all samples, one archaeal and 21 bacterial phyla were identified. The ten most abundant phyla recruited >98% of the reads generated from the microbial communities of both the control and *A. taxiformis* amended vessels (Figure 4). Microbiomes throughout the experiment, regardless of experimental condition or time, were dominated by *Bacteroidetes*, *Firmicutes*, and *Proteobacteria*. The *Bacteroidetes:Firmicutes* ratio decreased in both conditions over the course of the experiment, suggesting influence due to the experimental system (Figure 4). With the drastic decrease in CH_4_ in mind, the differences between the two groups were investigated at a finer resolution by exploring the abundance dynamics of the Archaeal phylum Euryarchaeota, which include the methanogenic Archaea. Based on the 16S rRNA gene profiles, five genera of methanogenic Archaea were identified in all stages of the experiment. The five genera: *Methanobrevibacter*, *Methanosphaera*, *vadin CA11* of the *Methanomassiliicoccacaea* family, *Methanoplanus* and *Methanimicrococcus* accounted for all reads recruited by the Euryarchaeota. *Methanobrevibacter* and *Methanosphaera* accounted for >99% of the reads assigned to methanogens. While CH_4_ production decreased in the *A. taxiformis* amended vessels 12 hr after the first feeding event, abundance of methanogenic Archaea in the two conditions did not differ significantly at individual time points (Figure 5). However, the average relative abundance of Euryarchaeota over the duration of the experiment were lower in the *A. taxiformis* amended vessels compared to control vessels (1.38 and 1.79% respectively, p ≤ 0.03).

**Figure 4.**
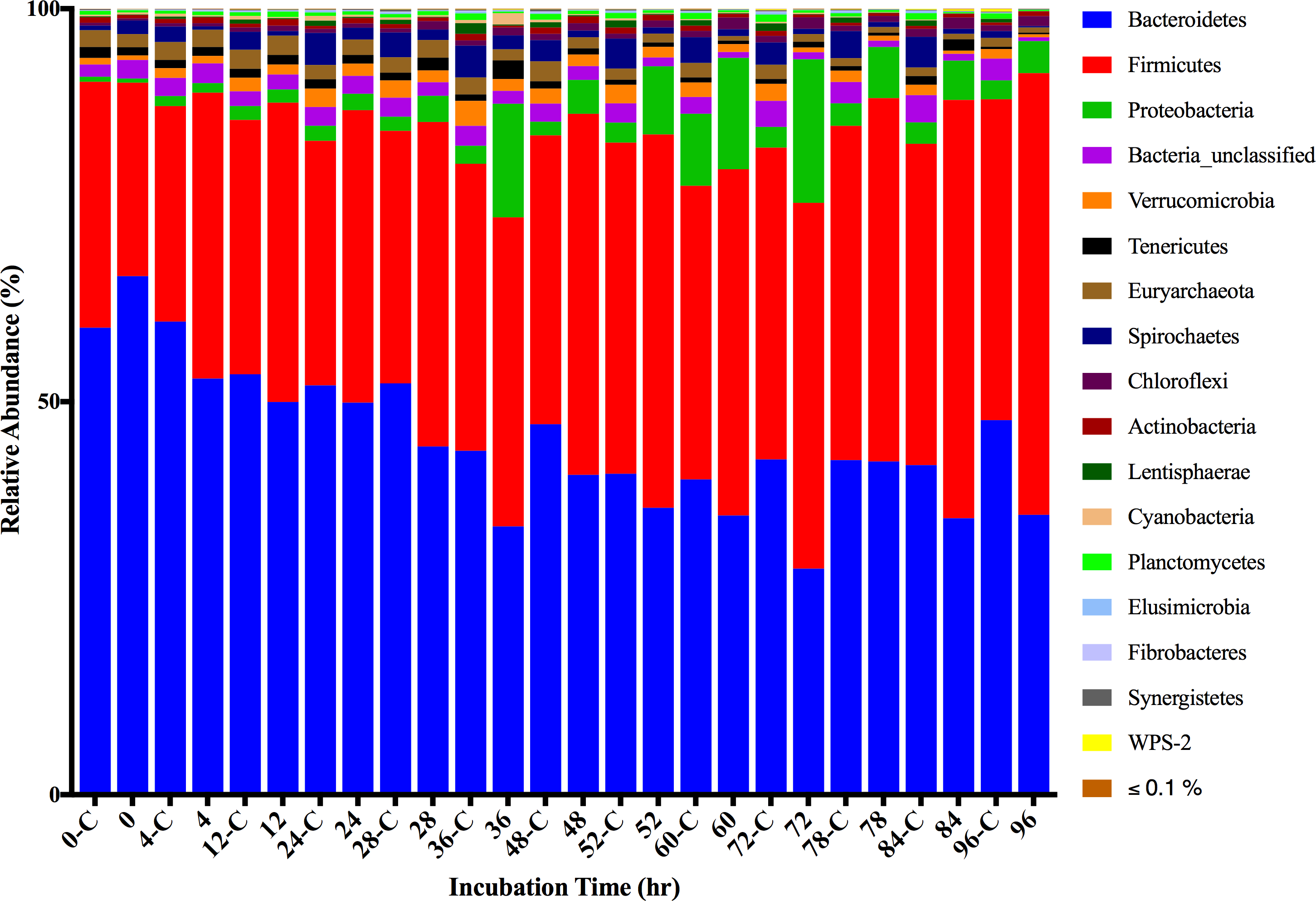
Relative Abundance of Phyla during *in-vitro* fermentation. Fermentations were performed in three *in-vitro* vessels (n=3). Incubation times annotated with “C” represent control conditions.

**Figure 5.**
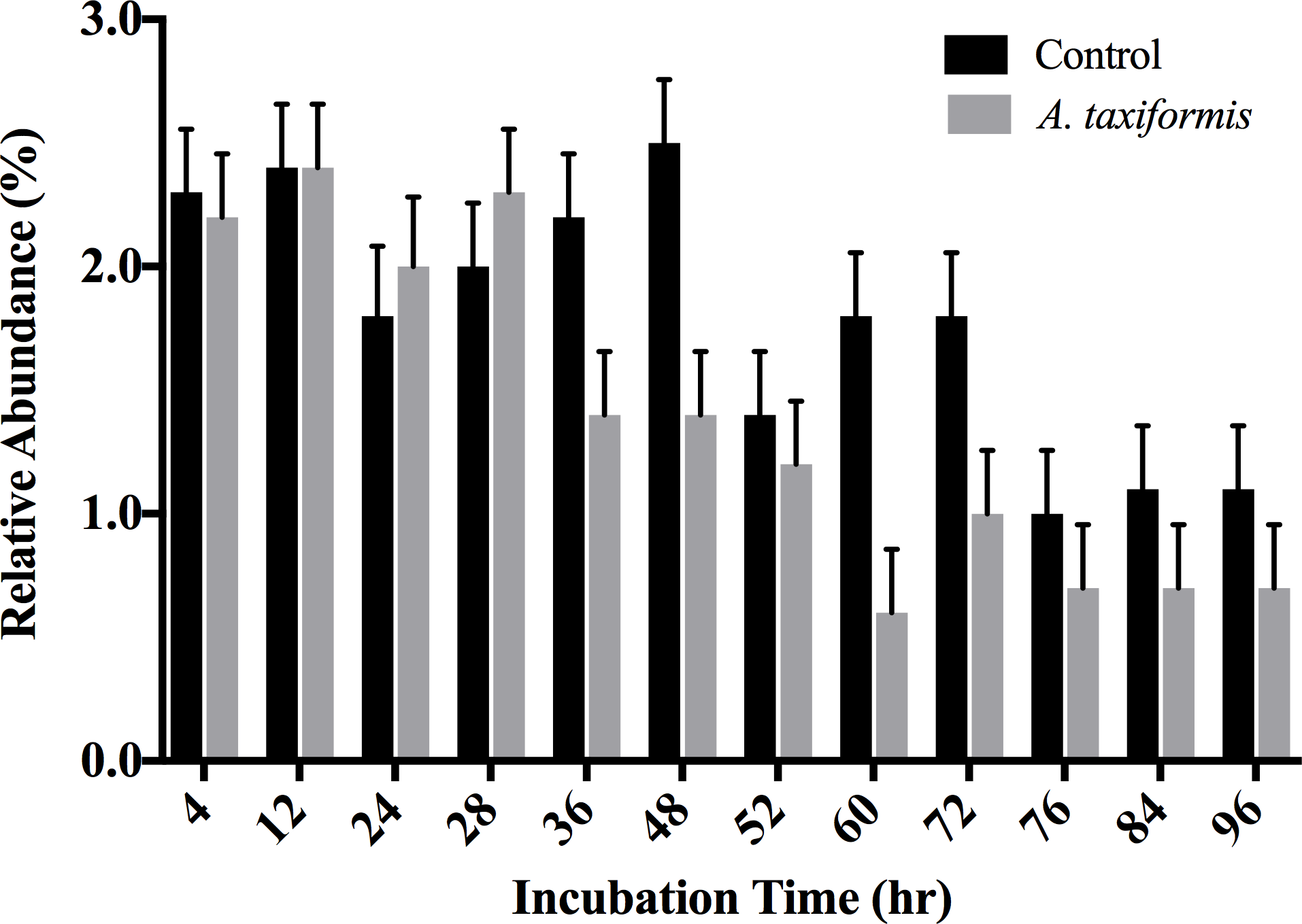
Relative abundance of Euryarchaeota during in-vitro fermentation. Fermentations were performed in three in-vitro vessels (n=3). Error bars indicate standard error of the mean.

## DISCUSSION

A significant reduction in CH_4_ production was found when evaluating the effects of *A. taxiformis* on ruminal fermentation characteristics *in-vitro* at a 5% OM inclusion rate. Results from the overall experiment show an approximate decrease in TGP by ~50% and in CH_4_ production by ~95%, which is similar to the reduction that was reported previously when the effect of *A. taxiformis* on CH_4_ production from beef cattle was investigated [10, 18, 30] Carbon dioxide production remained similar between the control and *A. taxiformis* amended vessels. Comparison of total and individual VFA between vessels did not suggest any difference in VFA production at any specific time point with the 5% OM inclusion rate. A significant reduction of CH_4_ was measured 12 hrs after *A. taxiformis* amendment (Figure 1), while CO_2_ production and VFAs profiles remained unchanged throughout the fermentation process (Figures 1 & 2). This suggests that the amendment of SBR supplemented with *A. taxiformis*, inhibits methanogenesis but not necessarily overall microbial growth, *per se*. This targeted effect on a specific metabolic function and hence a functional group within the microbiome was also elucidated from the 16S rRNA profiles of the *in-vitro* rumen system. Whereas the overall assemblages of the microbiome associated with the treatment and control fermentation vessels of the *in-vitro* rumen system remained rather similar throughout the duration of the fermentation process (Figure 4), changes in the relative abundance of members belonging to the Euryarchaeota, the taxonomic group that encompasses the main rumen methanogens, could be observed as early as 36 hr after the initiation of the experiment. Although a semi-continuous batch fermentation system, as utilized for this study, is capable of maintaining more rumen like conditions, mainly through maintaining adequate pH and nutrient levels, when compared to a simple batch fermentation process, a wash-out of the more sensitive rumen microbes (i.e. protozoa) is inevitable [31]. It is well known that there is a mutualistic relationship between protozoa and methanogens [32, 33], and it has been shown before that the removal of rumen protozoa results in a reduction of the methanogen population and methanogenesis during enteric fermentation [34, 35]. Hence, the decrease in relative abundance of Euryarchaeota observed for the control vessels at later time points of the experiment is most likely an artifact caused by the inability of the *in-vitro* systems to maintain protists over an extended period of time.

### Propionate:acetate ratio increased in treatment vessels

Over the course of the experiment, the propionate:acetate ratio increased (*p* < 0.001) in treatment vs control groups. The first step of the formation of acetate in the rumen releases metabolic hydrogen which acts as a hydrogen donor to methanogenic archaea and therefore facilitates the production of CH_4_ in the rumen [36]. In contrast, propionate acts as a competing hydrogen sink [4, 37]. The increased propionate:acetate ratio suggest that hydrogen is, at least in some part, being redistributed to propionate, which may help explain a portion of the methane reduction seen here. In the context of dairy cattle and milk production, the increased propionate:acetate ratio seen in vessels amended with *A. taxiformis* may forecast an altered milk composition *in-vivo*. A decreased propionate:acetate ratio is associated with increased milk fat, and total milk yield is positively associated with butyrate and propionate in the rumen [38]. Under this paradigm, *A. taxiformis* supplementation has the potential to increase total milk yield, however may also negatively impact milk fat content.

### Microbial communities overcame the stress of treatment

We observed that seaweed amendment has effects consistent with the Anna Karenina Hypothesis, which posits that disturbances act to increase differentiation of microbial communities [39]. Specifically, we found that communities in treatment vessels differentiated increasingly from each other up to hour 72, after which they re-converged (Figure 3A). Hence, the rumen microbial community undergoes changes that are both slow and variable in response to seaweed amendments. However, these changes do not appear to be associated with variability in reduction of gas production. While seaweed amendment may pose an initial stress on the rumen microbial community, measured by the increased differentiation between treatment vessels, after only 72 hours under recurrent daily stress (feeding), the β-diversity between communities in amended vessels stabilized.

### *A. taxiformis* is a potential mineral supplement

Nutritional analysis of *A. taxiformis* revealed that the red macroalgae has high levels of important minerals including calcium, sodium, iron, and manganese (Table 2) suggesting that in addition to its methane reduction potential, *A. taxiformis* may also be used to increase mineral availability to basic rations. *In-vivo* studies directed towards monitoring mineral transfer from feed into product should be conducted next to facilitate a better understanding of whether or not minerals or other compounds present in seaweed can be found in milk or meat of the consuming animals. While halogen compounds have been reported as important players in the bioactive process of methane reduction, previous studies using seaweed as a feed supplement found that iodine, which is abundant in brown algae, is found in the milk of cows to which it is fed [40].

## CONCLUSIONS

The methane reducing effect of *A. taxiformis* during rumen fermentation of feedstuff widely used in California, makes this macroalgae a promising candidate as a biotic methane mitigation strategy in the largest milk producing state in the US. The organic matter inclusion required to achieve such a drastic decrease in methane is low enough to be practically incorporated in the rations of average dairy operations. Significant limitations to the implementation of *A. taxiformis* and potentially other algae include the infrastructure and capital necessary to make these products commercially available and affordable. Furthermore, our understanding of the host microbe interactions during seaweed amendment are limited. In order to obtain a holistic understanding of the biochemistry responsible for the significant reduction of methane, and its potential long-term impact on ruminants, gene expression profiles of the rumen microbiome and the host animal are warranted.

## ABBREVIATIONS

16S rRNA: 16 Svedberg ribosomal Ribonucleic Acid
AMOVA: Analysis of Molecular Variance
bp: base pair
C: Celsius
CH_4_: Methane
Co: Company
CO_2_: Carbon Dioxide
DM: Dry Matter
DNA: Deoxyribonucleic Acid
FID: Flame Ionization Detector
g: Gram
GC: Gas Chromatography
Hrs: Hours
IACUC: Institution of Animal Care and Use Committee
ml: Milliliters
OM: Organic Matter
OTU: Operational Taxonomic Unit
PCoA: Principal Coordinate Analysis
PCR: Polymerase Chain Reaction
PVC: Poly Vinyl Chloride
SBR: Super Basic Ration
SD: Standard Deviation
TDN: Total Digestible Nutrients
TGP: Total Gas Production
VFA: Volatile Fatty Acid

## METHODS

### Animals, diets and rumen content collection

All animal procedures were performed in accordance with the Institution of Animal Care and Use Committee (IACUC) at University of California, Davis under protocol number 19263. Rumen content was collected from two rumen fistulated cows, one Jersey and one Holstein, housed at the UC Davis Dairy Unit. Animals were fed a dry cow total mixed ration (50% wheat hay, 25% alfalfa hay/manger cleanings, 21.4% almond hulls, and 3.6% mineral pellet (Table 1). Three liters of rumen fluid and 60 g of rumen solids were collected 90 min after morning feeding. Rumen content was collected via transphonation using a perforated PVC pipe, 500 mL syringe, and Tygon tubing (Saint-Gobain North America, PA, USA). Fluid was strained through a colander and 4 layers of cheesecloth into two 4L pre-warmed, vacuum insulated containers and transported to the laboratory.

### *In-vitro* feed and feed additive composition and collection

Due to its wide utilization in the dairy industry for cows during lactation, super basic ration (SBR) was used as feed in the *in-vitro* experiment. SBR was composed of 70% alfalfa pellets, 15% rolled corn, and 15% dried distillers’ grains (Table 3). Individual components were dried at 55°C for 72 hours, ground through a 2 mm Wiley Mill (Thomas Scientific, Swedesboro, NJ) and manually mixed. *Asparagopsis taxiformis* used as feed additive was provided in kind from the Commonwealth Scientific and Industrial Research Organization (CSIRO) Australia. The macroalgae was in its filamentous gametophyte phase when collected near Humpy Island, Keppel Bay, QLD (23°13’01”S, 150°54’01”E) by MACRO (Center for Macroalgal Resources and Biotechnology) of James Cook University (JCU) in Townsville, QLD. The collected biomass was frozen and stored at -15°C then shipped to Forager Food Co. in Red Hills, Tasmania, AUS, where it was freeze dried and milled (2-3 mm) to ensure a uniform product. Chemical composition of SBR and of *A. taxiformis* were analyzed at Cumberland Analytical Services (Waynesboro, PA).

**Table 3.**
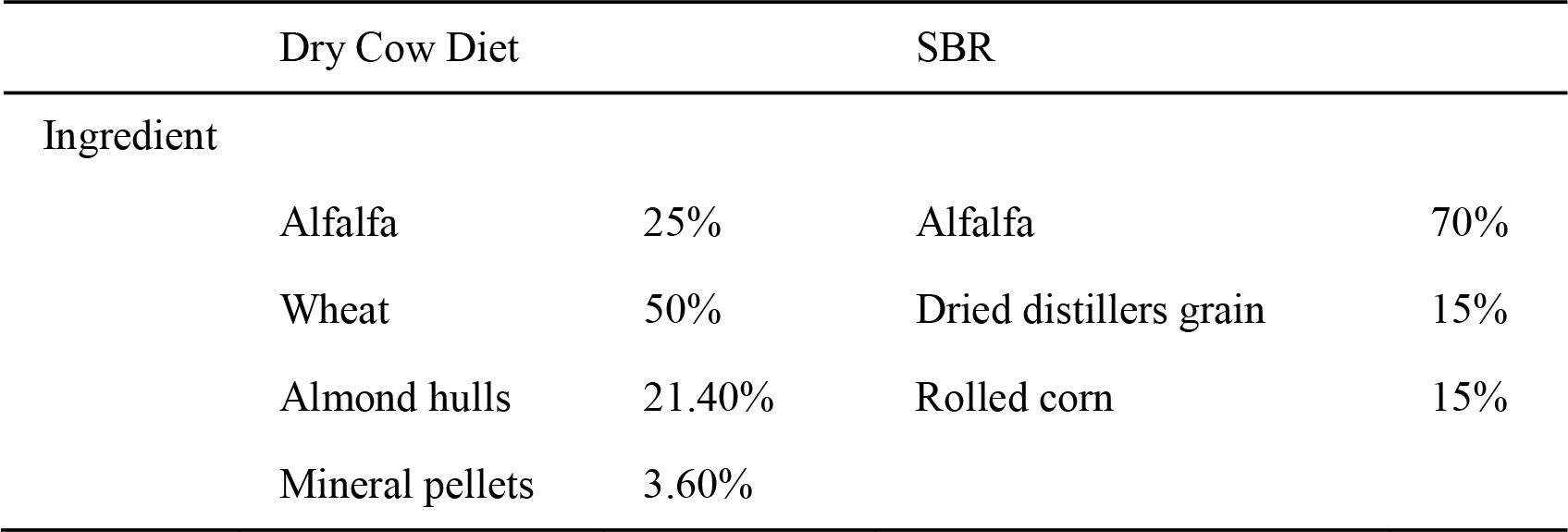
Composition of dry cow diet and super basic ration (SBR).

| Dry Cow Diet |  | SBR |  |
| --- | --- | --- | --- |
| Ingredient |  |  |  |
| Alfalfa | 25% | Alfalfa | 70% |
| Wheat | 50% | Dried distillers grain | 15% |
| Almond hulls | 21.40% | Rolled corn | 15% |
| Mineral pellets | 3.60% |  |  |

### Engineered (*in-vitro*) rumen system

An advanced semi-continuous fermentation system, with six 1L vessels with peristaltic agitation, based on the rumen simulation technique (RUSITEC) developed by Czerkawski and Breckenridge [41] was used to simulate the rumen in the laboratory.

### Experimental design

*Equilibration* (Day 0): Temperature, pH and conductivity of the rumen fluid and solids were recorded using a mobile probe (Extech Instruments, Nashua, NH). Rumen fluid, 3L, from each cow were combined with 2L of artificial saliva buffer [42] homogenized and then split into two 3L aliquots. Rumen solids, 15 g, from each animal were sealed in Ankom concentration bags (Ankom, Macedon, NY) and added to each equilibration vessel (30g of rumen solids per vessel total). Three concentrate bags containing 10g of SBR each were added to each vessel. One of the vessels was also inoculated with 1.5 g of *A. taxiformis* to equilibrate microbial populations to the treatment prior to the start of the experiment. SBR was ground in a 2 mm Wiley Mill before being added to each concentrate bag to increase substrate availability and therefore producing similar particle sizes that which the mastication function *in-vivo* provides to the animal. The two vessels were then placed in a 39°C water bath and stirred with a magnetic stir bar for a 24 hour equilibration period.

*Fermentation* (Days 1-4): After 24 hours of equilibration, temperature, pH, and conductivity of the rumen fluid were recorded to determine stability of the vessels and their content. Each of the 6 *in-vitro* rumen vessels were randomly designated as either treatment or control vessel and filled with 750 mL of the corresponding fluid from the equilibration vessels. Location of the vessels within the *in-vitro* platform were randomly allocated.

Each vessel received one concentrate bag of SBR from its respective equilibration vessel and one new concentrate bag. Control concentrate bags contained 10 g SBR. Treatment concentrate bags contained 10 g SBR plus 5% (OM) *A. taxiformis*. To simulate rumen retention time, each of the feedbags were incubated in the allocated fermentation vessel for 48 hours. Temperature, pH, and conductivity were measured every 24 hours prior to exchanging one of the concentrate bags (feeding). After each feeding, all vessels were flushed with N_2_ to maintain anaerobic conditions within the reactors. Individual reactor vessels of the artificial rumen system were connected to a reservoir containing artificial saliva buffer. A peristaltic pump delivered 0.39 mL/min of buffer to each vessel throughout the course of the experiment. Gas bags (Restek, USA) and overflow vessel were used to continuously collect generated gas and effluent fluid. Effluent vessels were chilled with ice to mitigate residual microbial activity.

### Sample collection and analysis

Liquid and gas sample collections took place at 3 time points every 24 hours for 4 days. Time point intervals were 4, 12, and 24 hours post-feeding each day. Fluid samples were collected in mL tubes, flash frozen in liquid nitrogen, and stored at -20°C until processed. Gas bags were collected at each time series interval for analysis of total gas production, CO_2_ and CH_4_ concentrations. Gas volume was measured with a milligas flow meter (Ritter, Germany) by manual expulsion of the collection bag.

### Volatile fatty acid and greenhouse gas analysis

To determine VFA profiles, Gas Chromatography-Flame Ionization detection (GC-FID) was used. Fermentation fluid was prepared for VFA analysis by mixing with 1/5th volume 25 % metaphosphoric acid, and centrifugation. Supernatant was filtered through a 0.22 µm filter and stored in amber autosampler vials at 4 °C until analysis. The GC conditions were as follows: analytical column RESTEK Rxi^®^ – 5 ms (30 m × 0.25 mm I.D. × 0.25 µm) film thickness; the oven temperature was set to 80°C for 0.50 min, and followed by a 20°C/min ramp rate until 200°C, holding the final temperature for 2 min; carrier gas was high purity helium at a flow rate of 2.0 mL/min, and the FID was held at 250°C. A 1 µL sample was injected through Split/Splitless Injectors (SSL), with an injector base temperature set at 250°C. Split flow and split ratio were programmed at 200 and 100 mL/min respectively. To develop calibration curves, certified reference standards (RESTEK, Bellefonte, PA) were used. All analyses were performed using a Thermo TriPlus Autosampler and Thermo Trace GC Ultra (Thermo Electron Corporation, Rodano Milan, Italy).

Methane and CO_2_ were measured using an SRI Gas Chromatograph (8610C, SRI, Torrance, CA) fitted with a 3’x1/8” stainless steel Haysep D column and a flame ionization detector with methanizer (FID-met). The oven temperature was held at 90°C for 5 minutes. Carrier gas was high purity hydrogen at a flow rate of 30 ml/min. The FID was held at 300°C. A 1 mL sample was injected directly onto the column. Calibration curves were developed with an Airgas certified CH_4_ and CO_2_ standard (Airgas, USA).

### DNA extraction

DNA extraction was performed using the FastDNA SPIN Kit for Soil (MP Biomedicals, Solon, OH) with ~500 mg of sample according to the manufacturer’s protocol. DNA was subsequently purified with a Monarch^®^ PCR & DNA Cleanup Kit (New England Biolabs, Ipswich, MA) following the manufacturer's instructions. Extracted DNA was stored at -20°C until subsequent PCR amplification and amplicon sequencing.

### PCR amplification, library preparation, and sequencing

The V4-V5 hypervariable region of the 16S rRNA gene was sequenced on Illumina’s MiSeq platform using the 515yF (3’-GTG YCA GCM GCC GCG GTA A-5’) and 926pfR (3’-CCG YCA ATT YMT TTR AGT TT-5’) primer pair (Research and Testing, Lubock Texas; [43, 44] For sequencing, forward and reverse sequencing oligonucleotides were designed to contain a unique 8 nt barcode (N), a primer pad (underlined), a linker sequence (italicized), and the Illumina adaptor sequences (bold).

Forward primer: **AATGATACGGCGACCACCGAGATCTACAC-**NNNNNNNN-<u>TATGGTAATT</u>-*GT-*GTGYCAGCMGCCGCGGTAA;

Reverse primer: **CAAGCAGAAGACGGCATACGAGAT-**NNNNNNNN-<u>AGTCAGTCAG</u>-*GG-*CCGYCAATTYMTTTRAGTTT.

Barcode combinations for each sample are provided in Supplementary Table S4. Each PCR reaction contained 1 Unit Kapa2G Robust Hot Start Polymerase (Kapa Biosystems, Boston, MA), 1.5 mM MgCl_2_, 10 pmol of each primer, and 1μL of DNA. The PCR was performed using the following conditions: 95°C for 2 min, followed by 30 cycles at 95°C for 10 s, 55°C for 15 s, 72°C for 15 s and a final extension step at 72°C for 3 min. Amplicons were quantified using a Qubit instrument with the Qubit High Sensitivity DNA kit (Invitrogen, Carlsbad, CA). Individual amplicon libraries were pooled, cleaned with Ampure XP beads (Beckman Coulter, Brea, CA), and sequenced using a 300 bp paired-end method on an Illumina MiSeq at RTL Genomics in Lubbock Texas. Raw sequence reads were submitted to NCBI’s Sequence Read Archive under the SRA ID: SRP152555.

### Sequence analysis

Sequencing resulted in a total of 1,251,439 raw reads, which were analyzed using mothur v1.39.5 [45] using the MiSeq SOP accessed on 3/10/2018 [46]. Using the *make.contigs* command, raw sequences were combined into contigs, which were filtered using *screen.seqs* to remove sequences that were >420 bp or contained ambiguous base calls to reduce PCR and sequencing error. Duplicate sequences were merged with *unique.seqs*, and the resulting unique sequences were aligned to the V4-V5 region of the SILVA SEED alignment reference v123 [47] using *align.seqs*. Sequences were removed if they contained homopolymers longer than 8 bp or did not align to the correct region in the SILVA SEED alignment reference using *screen.seqs*. To further denoise the data, sequences were pre-clustered within each sample allowing a maximum of 3 base pair differences between sequences using *pre.cluster.* Finally, chimeric sequences were removed using VSEARCH [48].

Quality filtered sequences were grouped into OTUs based on 97% sequence identity and classified using the Bayesian classifier and the Greengenes database (August 2013 release of gg_13_8_99) [49] with *classify.seqs*. Sequences that classified as mitochondria, chloroplasts, eukaryotic, or of unknown origin were removed using *remove.lineage.* Samples were rarefied to 6,467 sequences per sample, the smallest number of sequences across all collected samples. Singleton abundances were calculated with *filter.shared.* Chao1 diversity [50], Good’s coverage [51], Shannon [52], and inverse Simpson indices were calculated using *summary.single* to quantify coverage and α-diversity.

### Alpha-diversity

To estimate the microbial diversity within each group, first, rarefaction analyses were performed (Supplementary Figures S1) and species richness and diversity indices were calculated (Supplementary Table S2.). Variance of the microbial community between and among the different vessels were quantified using a θ_YC_ distance matrix [53].

### Beta-diversity

To investigate slow-acting effects of seaweed addition on microbiome communities, we computed Bray-Curtis dissimilarity (β-diversity) [54] between pairs of samples, both within vessels at different time points, and between vessels at identical time points. We also considered Jaccard dissimilarity which only reflects community composition and not relative abundance, but found similar results and so only report the results for Bray-Curtis dissimilarity. We independently computed β-diversity at the genus, family, order, class, and phylum level to assess whether the patterns that we found were dependent on taxonomic resolution. All analyses were performed using custom-written Java, SQL, and Bash code available at https://github.com/jladau.

### Statistical analysis

Analysis of molecular variance (AMOVA) [55] was used to identify significant differences in community structure between treatment and control vessels using a θ_YC_ distance matrix for the *amova* command in Mothur. The complete results of these statistical tests between each time interval combination is included in the supplementary data.

Gas, VFA, and Euryarchaeota abundance data were analyzed using the linear mixed-effects model (lme) procedure using the R statistical software (version 3.1.1) [56]. The statistical model included treatment, day, time point, treatment×day×time point interactions, treatment×day interactions, treatment×time point interactions, day×time point interactions and the covariate term, with the error term assumed to be normally distributed with mean = 0 and constant variance. Orthogonal contrasts were used to evaluate treatments vs. control, linear, and quadratic effects of treatments. Significant differences among treatments were declared at p ≤ 0.05. Differences at 0.05 < p ≤ 0.10 were considered as trend towards significance.

## ETHICS APPROVAL

All animal procedures were performed in accordance with the Institution of Animal Care and Use Committee (IACUC) at University of California, Davis under protocol number 19263.

## CONSENT FOR PUBLICATION

Not applicable

## AVAILABILITY OF DATA AND MATERIAL

Sequence data generated during this study are available through NCBI’s Sequence Read Archive under the SRA ID SRP152555. Custom-written Java, SQL, and Bash code is available at https://github.com/jladau. All other data is included in this published article and its supplementary information files.

## AUTHORS’ CONTRIBUTIONS

Designed the experiment: BRoque, CBrooke, EKebreab, JSalwen and MHess; Performed the experiments: BRoque, CBrooke, MHess and NNajafi; Generated and analyzed the microbiome data: BRoque, CBrooke, EEloe-Fadrosh, JLadau, MHess and NNajafi. Generated and analyzed GC data: BRoque, CBrooke, LMarsh, LSingh, MHess, NNajafi, PPandey; Wrote the paper: BRoque, CBrooke, EEloe-Fadrosh, EKebreab JLadau, JSalwen LMarsh, MHess and TPolley.

## ACKNOWLEDGEMENTS

The authors would like to thank Kyra Smart, Susan Parkyn and Ania Kossakowski for their assistance in maintaining the artificial rumen system.

## SUPPLEMENTARY INFORMATION

Supplementary information is available at BMC Microbiomes’ website.

Supplemental File: A.tax_supplemental_Tables_Figures_20180711

Supplementary Table S1. Quality filtering and OTU distribution at each incubation time.

Supplementary Table S2. Diversity indices at each incubation time.

Supplementary Figures S1A, S1B, S1C. Rarefaction curves of equilibration, control and *A. taxiformis* amended vessels respectively.

Supplementary Figure S2. Principle Coordinate Analysis plot.

Supplementary Table S3. OTU table.

Supplementary Table S4. Raw sequence barcodes for archived 16S rRNA gene amplicon data.

Supplementary Table S5. Results of AMOVA and HOMOVA statistical tests.

